# IgG4 antibody repertoire diversity is stable over several years in health and autoimmune disease

**DOI:** 10.64898/2026.09.30.755566

**Authors:** Linus Wollenweber, Laurent M. Paardekooper, Yvonne J.M. Campman, Jessica M. van Bokkum, Martijn R. Tannemaat, Jan J.G.M. Verschuuren, Arjan Barendregt, Albert J.R. Heck, Albert Bondt, Maartje G. Huijbers

## Abstract

Despite its relative anti-inflammatory nature, IgG4 is involved in a variety of pathological conditions such as autoimmune disease and cancer. To investigate whether the IgG4 repertoire presents any abnormalities in diseases where IgG4 autoantibodies predominate, we charted longitudinal IgG4 repertoires in health, in an IgG4-mediated autoimmune disease (AID; muscle-specific kinase myasthenia gravis (MuSK MG)), and in IgG1/IgG3 mediated AIDs (acetylcholine receptor (AChR) MG and Lambert-Eaton myasthenic syndrome (LEMS). In total, we assessed 218 samples of 69 donors using mass spectrometry-based antibody repertoire analysis, covering a timespan of up to eight years.

Despite being superficially inconspicuous in clonal diversity and abundances, we detected a slightly elevated number of IgG4 clones in MuSK MG patients compared to healthy donors. Notably, this observation did not coincide with drastic repertoire changes during treatment, suggesting that a large portion of the IgG4 repertoire is not significantly affected by common MuSK MG treatment regimens.

Comparing the IgG4 antibody repertoires of all donors, we found that they were exceptionally unique for each individual. Against what would be expected in the case of a constantly adapting antibody repertoire, a large portion of the IgG4 clones remained stable without any further maturation. This observation even extended to patients receiving B cell depleting or IgG modulating treatments. Our findings suggest that a dominating fraction of the IgG4 pool may be produced by long-lived plasma cells, which are unaffected by disease or medical interventions.

## Introduction

Antibodies form a core defense mechanism against pathogens^1^, allergens^2^, and toxins^3^. The different human antibody (sub)classes (IgM, IgD, IgG1-4, IgA1-2 and IgE) are distinguished by their heavy chain constant regions (Fc), which give rise to differences in tissue distribution^4^, immune complex formation^5^, complement activation^6^, and effector cell interactions through Fc receptors^7^.

Of all IgG subclasses, IgG4 is often regarded as the odd one out in both structure and function^5, 8^. Structurally, IgG4 is uniquely capable of forming *intra*-heavy chain disulfide bonds in the hinge region. This allows one bivalent IgG4 molecule to dissociate into two half-molecules that can exchange with other IgG4 half molecules to form functionally monovalent, bispecific antibodies^9^ (Figure 1A). This stochastic process is called Fab-arm exchange (FAE) and affects all endogenous serum IgG4s^10^. Functionally, IgG4 interacts only weakly with Fcγ receptors and complement factors, resulting in an anti-inflammatory phenotype^7, 11-13^.

**Figure 1:**
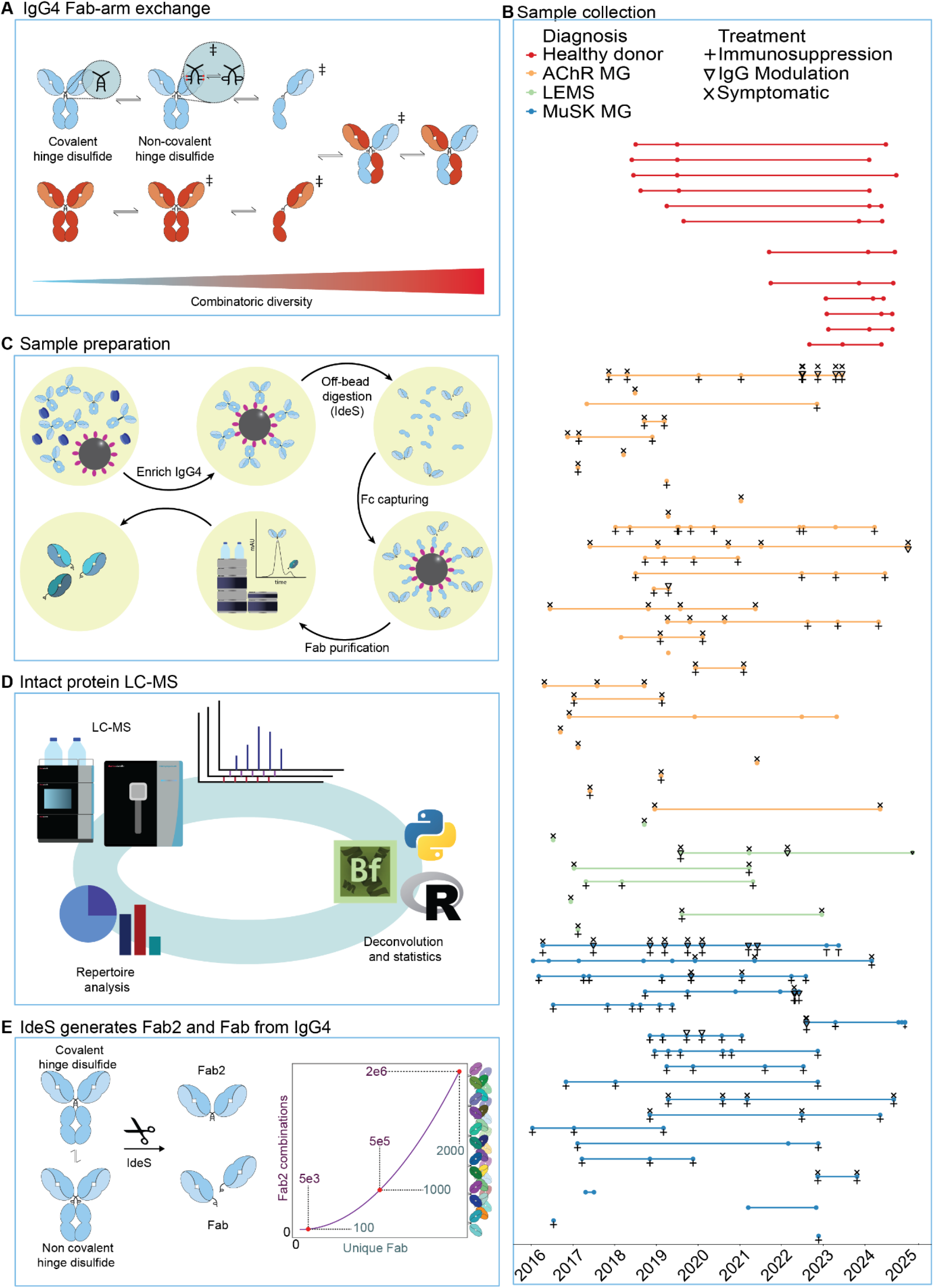
Overview of IgG4 repertoire analysis and donor cohort. **(A)** IgG4 stands out from other IgG classes by its ability to dissociate and undergo FAE. Cartoon adapted from Wollenweber *et al*.^10^ **(B)** A cohort of 69 donors was sampled longitudinally, totaling 218 serum samples. Healthy IgG4 repertoires from 12 donors (red) were compared to IgG4 repertoires from patients diagnosed with AChR myasthenia gravis (orange, n = 29), MuSK myasthenia gravis (blue, n = 20), and LEMS (green, n = 8). Sampling points are indicated as dots, and treatments are grouped into immunosuppressive (plus sign), IgG modulating (triangle sign), and symptomatic treatments (cross sign). **(C)** IgG4 antibodies were purified from serum using affinity matrices. The IdeS protease was used to release Fab2 and Fab from the antibodies. Fc regions were removed via affinity capture, and Fab fragments were separated from Fab2 using size exclusion chromatography (SEC). **(D)** Fabs were mass analyzed by protein-centric LC-MS, and masses were retrieved using BioPharmaFinder with additional downstream analysis using Python and R. **(E)** As a consequence of the hinge disulfide equilibrium in IgG4, IdeS digestion yields two products: Fab and Fab2. Only Fab are analytically accessible as the complexity of Fab2s is massively inflated. Cartoon adapted from Wollenweber *et al*.^10^.

Even though IgG4 is the least abundant IgG subclass^14^, it predominates specific responses that are associated with chronic or repeated antigen exposure^15^, such as allergy^16-18^, parasitic infection^19^, or autoimmunity^20-22^. While most antibody-mediated autoimmune diseases (AIDs) are driven by the pro-inflammatory IgG1 and IgG3 subclasses^23^, a subset of AIDs is strongly linked to pathogenic IgG4 responses^20, 24^. For several IgG4-AIDs there is evidence for a direct link between FAE and pathogenicity^22-24^, of which muscle-specific kinase myasthenia gravis (MuSK MG) is an archetypical example^25^. Interestingly, other forms of myasthenic diseases are predominantly driven by IgG1- and IgG3-antibodies. In those diseases, autoantibodies are directed either against the acetylcholine receptor (AChR MG)^26^ or against voltage-gated calcium channels (VGCC) in Lambert-Eaton Myasthenic Syndrome (LEMS)^27^. Despite these mechanistic differences, all three conditions present with fatigable skeletal muscle weakness. Given the chronic nature of the IgG4 autoimmune response in MuSK MG, we hypothesized that their serum clonal IgG4 repertoires may be dominated by MuSK autoantibodies and be less diverse. To investigate whether IgG4 repertoire features in IgG4-AID are different from IgG1/IgG3-driven AIDs, we charted IgG4 repertoires across MuSK MG, AChR MG, LEMS, and healthy individuals.

Recent innovations in mass spectrometry allow sensitive and accurate analysis of intact antibody fragments and facilitate quantitative repertoire analyses with clonal resolution^10, 28, 29^. In the IgG4 antibody repertoire analysis approaches, IgG4 molecules are enriched from serum and subsequently cleaved using the hinge-directed protease IdeS, which releases the Fab domains that contain the complementarity determining regions (CDR) which determine the antibody target but also the uniqueness of clones^10^. Following clean-up, LC-MS detects each Fab (and thereby each IgG4 clone) at its unique mass and chromatographic retention time, resolving individual clones across the repertoire. This allows for longitudinal tracking of clonal dynamics, thereby providing a comprehensive readout of the overall IgG4 repertoire.

Here, using IgG4-directed antibody repertoire analysis, we charted the total IgG4 repertoires of 218 individual serum samples obtained from 69 donors, incorporating 20 MuSK MG patients (an IgG4-AID), 29 AChR MG patients, 8 LEMS patients (both IgG1/3-AIDs), and 12 healthy donors. In doing so, we reveal disease-related and treatment-related changes in the IgG4 repertoire with unmatched molecular precision. Each donor presented a unique IgG4 repertoire, which remained extremely stable both in sequence and individual clonal concentrations for more than five years. This hints that these repertoires originate from long-lived B cells that are resistant towards mutation and constantly produce their corresponding antibodies. MuSK MG patients showed no aberrations in IgG4 repertoire diversity, though their repertoires were slightly more polyclonal. In addition, their disease severity seemed to correlate with the number of detectable clones. Surprisingly, the effect of immunosuppressive and IgG-modulating treatments on the repertoire-dominating IgG4 clones of all donor groups was found to be marginal.

## Results

### Study population

We analyzed sera from 69 donors, encompassing patients with the IgG4-associated AID MuSK MG (n= 20), the IgG1/IgG3-associated AIDs AChR MG (n = 29) and LEMS (n = 8), and healthy controls (n = 12). The patients were included between 2016 and 2025 at the Leiden University Medical Center (LUMC) Department of Neurology outpatient clinic and material was collected during a planned visit and stored in the neuromuscular diseases biobank after signing informed consent. Patient material was selected for this study based on a positive autoantibody titer for either AChR, MuSK or VGCC using validated commercial radioimmunoprecipitation or ELISA assays at the clinical chemical lab and prioritizing those who had at least two follow up samples available. Healthy donor samples were obtained from the Leiden universal donor service at LUMC. For most donors, longitudinal samples were collected, spanning a range of up to 8 years (Fig. 1B). The number of time points and the time between sampling dates varied between patients as this was a prospective biobank collection. Demographics of the study populations, including male:female ratio, disease duration and clinical scores were comparable between donor groups (Table S1).

### IgG4-specific Fab profiling using LC-MS in AID

Using a recently introduced mass spectrometry-based workflow for IgG4 antibody repertoire analysis^10^, we aimed to investigate differences in serum IgG4 characteristics and repertoires across all serum samples from the IgG4-AID patients, IgG1/3-AID patients, and the healthy donors. A schematic overview of sample preparation and intact protein LC-MS is shown in Fig. 1C-D. Briefly, IgG4 antibodies were affinity-purified from serum samples. IgG4 Fab2 and Fab molecules, which carry the clonotype determining CDRs, were then released using the hinge-directed protease IdeS. The presence of both Fab2 and Fab fragments reflects the equilibrium between intra- and inter-heavy chain hinge disulfides characteristic of IgG4 (Fig. 1E). These distinct moieties can be separated by size exclusion chromatography (SEC). A simultaneous UV readout indicates the protein content during the separation, thereby providing quantitative information on the proportion of hinge isoforms.

Fraction collection during the SEC procedure further allows to separate the Fab and Fab2 fragments. As demonstrated earlier, Fab2 fragments are analytically not accessible for determining clonality in IgG4 samples, due to the extreme combinatorial diversity of Fab:Fab pairings caused by Fab arm exchange^10^ (Fig. 1E). Therefore, only the purified Fab fragments were subjected to liquid chromatography–mass spectrometry (LC–MS) for clonal separation and detection. In downstream analysis, each Fab clone is assigned an ID consisting of retention time and molecular mass, which enables the matching and differentiation of clones. This exact mass is a direct result of the precise amino sequence of each clone, and consequently any mutation in the sequence will lead to a change in the measured mass.

First, we characterized the IgG4 samples in terms of their macro characteristics, including the proportion of Fabs and the total clone number. The proportion of Fabs in the IdeS-cleaved samples was determined by SEC-UV. Fabs originate from the subgroup of serum IgG4 molecules that, at the time of sample collection, consist of non-covalently bound half-molecules (i.e., the population capable of undergoing FAE at that moment). Across all samples, we found ∼22% Fab (group range 20.6-23.3% Fab, SD 3.7%) and ∼78% Fab2 (Fig. 2A). All observed differences were, although statistically significant, within the range of variation that would also be expected within healthy samples^10^.

**Figure 2:**
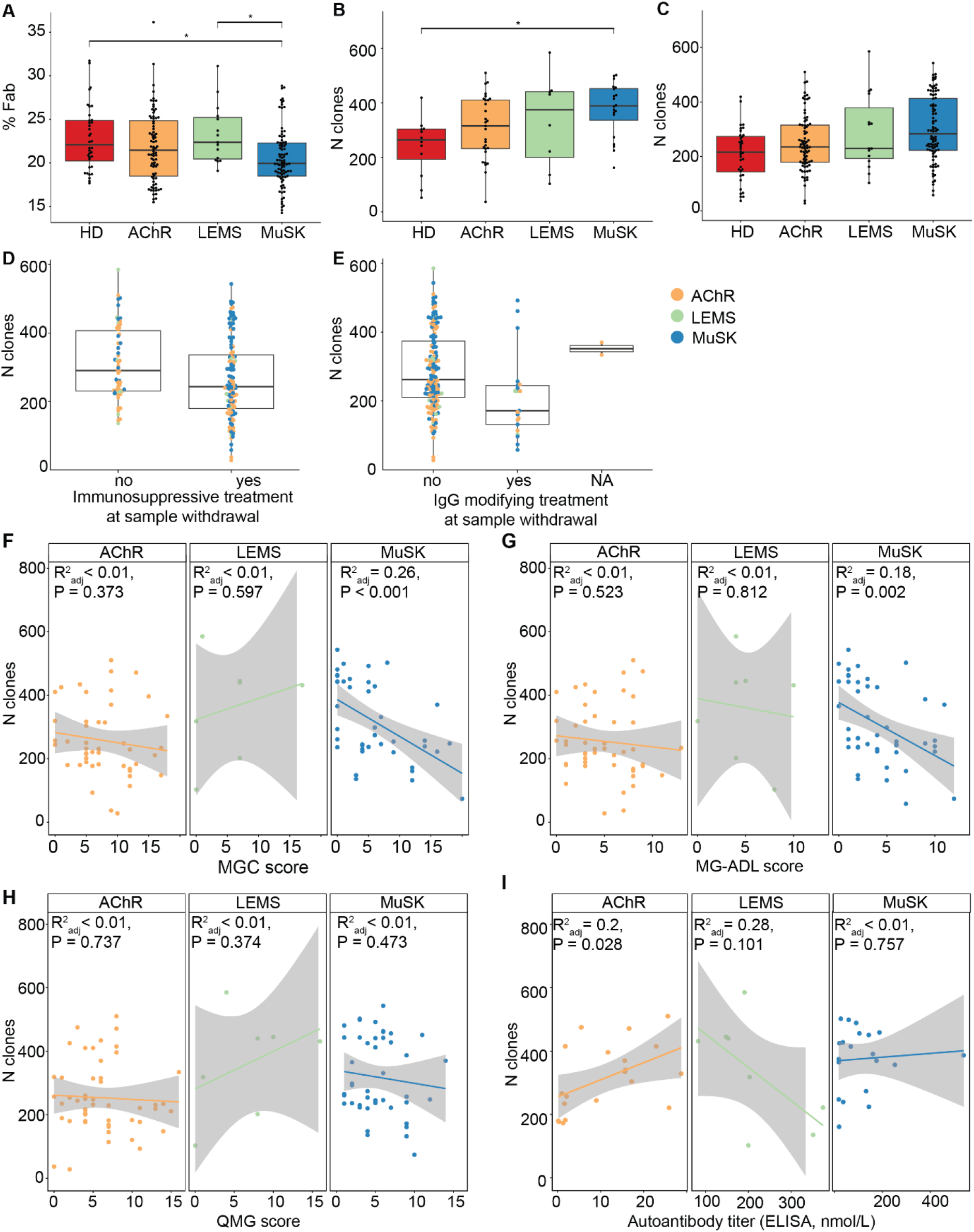
Structural capacity for FAE and clonality in IgG4 repertoires. **(A)** Proportion of IgG4 Fab in all samples, stratified by diagnosis. **(B, C)** IgG4 clone count detected at baseline (B) and in all samples (C), stratified by diagnosis. The color coding per physiological state is used throughout the figure. **(D, E)** IgG4 clone count detected in all samples, stratified by application of immunosuppressive treatment (D) and IgG modulating treatment (E). Patients treated with Rituximab were excluded from both analyses. **(F, G, H)** Linear regression analysis of IgG4 clone count detected at baseline with MGC disease severity score (F), MG-ADL disease severity score (G), and QMG disease severity score (H), stratified by diagnosis. **(I)** Linear regression analysis of IgG4 clone count detected at baseline with autoantibody titers matching the diagnosis (anti-AChR for AChR MG, anti-VGCC for LEMS, anti-MuSK for MuSK MG, respectively). Statistical analysis: * = p < 0.05, one-way ANOVA with Tukey’s HSD post-hoc test (panels A-C), linear mixed-effect model with disease duration and patien identifier added as random effects (panels D-E).

Next, we evaluated how many clones were detectable per sample. On average, we could detect 266 clones per sample, although there was notable variation (range 28-585). To exclude that changes in clone number detection would be an artifact of instrument drift over time, we repeatedly measured one sample at regular intervals as a technical control. This sample yielded on average 237 clones (SD ± 44) with high overlap in 6 replicates over the entire measurement period of 18 days (Sup. Fig. 1). Total IgG4 yields were estimated using the SEC data and found to be comparable between all donor groups (Sup. Fig. 2), although a non-significant subpopulation of MuSK MG samples showed increased IgG4 yields, and therefore likely also titers, matching a previous report^30^.

### MuSK MG patients have increased numbers of abundant IgG4 clones and these correlate with self-reported disease severity and treatment exposure

To assess IgG4 clonal diversity in the different AIDs, we compared the total number of detected unique IgG4 clones at baseline and found significantly more IgG4 clones in MuSK MG patients when compared to healthy donors (Fig 2B). Differences in clone count are no longer observed between groups when all timepoints are included in the analysis (Fig. 2C), which may be related to treatment effects. We therefore grouped treatments into three categories: immunosuppression (prednisolone, rituximab, complement inhibition, CellCept), IgG modulation (plasmapheresis, IVIg, efgartigimod), and symptomatic treatment (amifampridine, diaminopyridine, pyridostigmine). For the clonality analysis, we included only immunosuppressive or IgG modulating treatments, which affect either immune cells or antibodies, respectively. For both categories, we found no significant change in the number of detectable clones (Fig.2D, E). Due to rituximab being a large confounder, patients treated with rituximab were excluded from this analysis. Notably, most patients receive treatments from multiple categories at once, hampering detailed analysis of individual treatment effects. MuSK MG patients were significantly more likely to be treated with immunosuppression, while AChR MG patients received significantly more symptomatic treatments (Sup. Table 3).

To investigate whether the clone number (and thereby repertoire diversity) changes during the disease course, we correlated clone numbers to the main disease severity scoring instruments available for MG: QMG^31^, MGC^32^ and MG-ADL^33^ (Fig. 2F-H). For MuSK MG, we found significant negative correlations between the number of IgG4 clones in the sample and disease severity according to MGC and MG-ADL scoring. Interestingly, the MGC and MG-ADL scoring instruments are self-reported, whereas the QMG score is measured by a physician. For the IgG1-3-mediated diseases AChR MG and LEMS, no correlation was observed between the number of detected IgG4 clones and disease severity. This data suggests that overall IgG4 clonality remains stable independent of disease severity, except in MuSK MG. We investigated donor age as a possible confounder and found no correlation of age at sampling with number of clones detected (Sup. Fig. 3).

**Figure 3:**
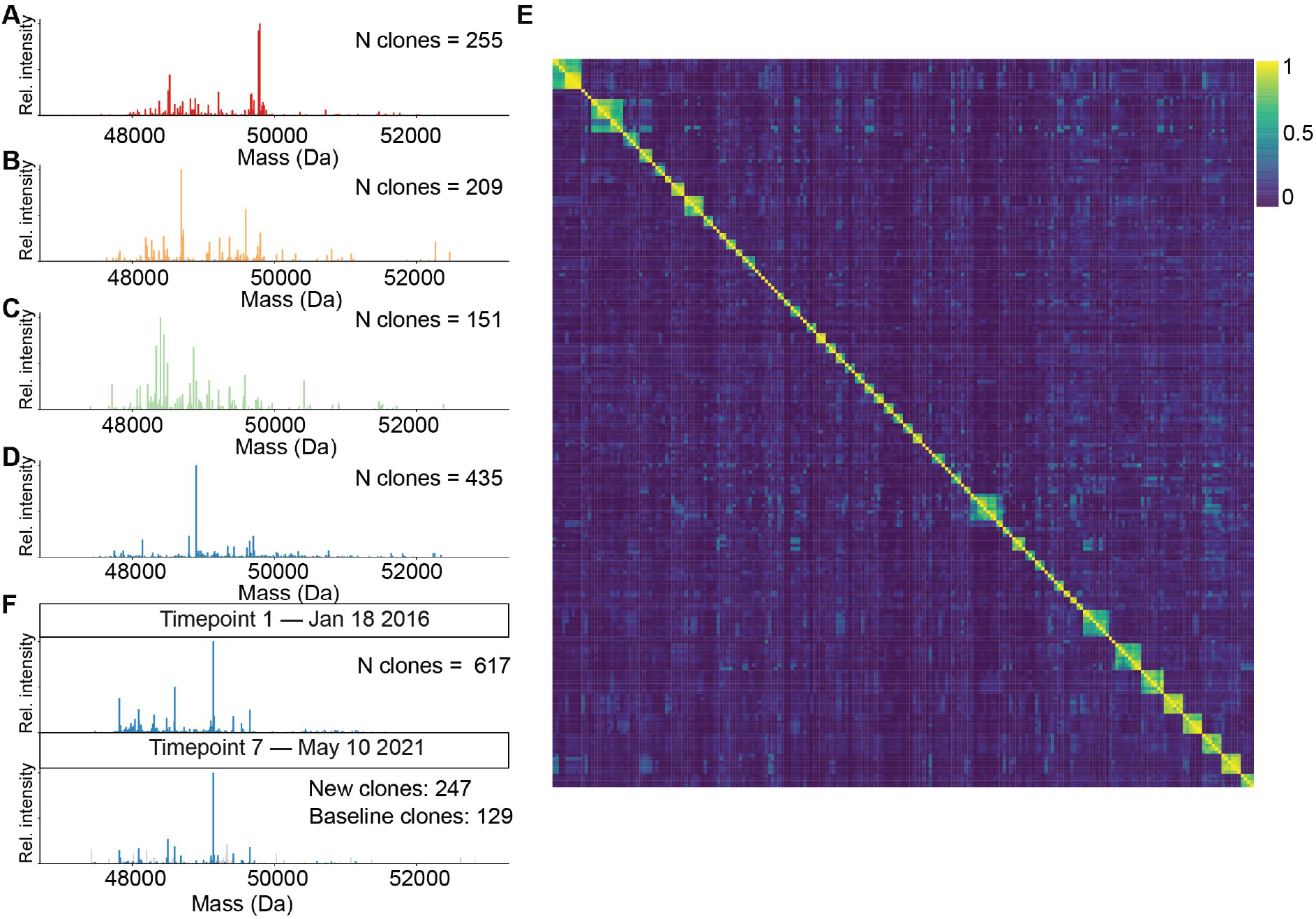
Cumulative overview of all IgG4 repertoires across 218 samples from 69 donors. **(A-D)** Mass plots of 4 representative IgG4 Fab clonal repertoires from various samples (healthy donor, AChR MG, LEMS and MuSK MG, respectively). Each bar represents a unique clone as recognized by matching mass and retention time. Clonal abundance is indicated by the height of the bars. **(E)** Pairwise intensity-based repertoire overlap between all recorded IgG4 repertoires. The similarity analysis provides a score indicating the similarity of each comparison. These scores range between 0 (no shared clonal intensity, dark blue) to 1 (all detected clonal intensity stems from clones in the counterpart, yellow). **(F)** IgG4 repertoires obtained for the same donor at baseline (top) and 5 years later (bottom). IgG4 clones also identified at baseline are colored; clones only identified after 5 years are highlighted in grey.

Lastly, we investigated correlations between baseline autoantibody titers determined by ELISA and number of IgG4 clones (Fig. 2I). We found a positive correlation between anti-AChR autoantibody titers and the total number of IgG4 clones. Notably, pathogenic autoantibodies in AChR MG are generally of the IgG1 and IgG3 subclasses. No correlation was found between anti-MuSK autoantibody titers and number of detected IgG4 clones. Due to limited sample numbers and missing data, attempts to investigate correlations in LEMS samples lack statistical confidence. In summary, our analyses revealed that MuSK patients have expanded IgG4 repertoires that correlate negatively with disease severity. This association is not explained by IgG4 autoantibody titers but may reflect the effects of immunotherapies on B cell survival and thus clonality.

### IgG4 repertoires are donor specific and remain unchanged for several years

Next, the IgG4 repertoire similarity between all individual samples was examined. This involved analyzing both the longitudinal repertoire stability within a single donor and any potential repertoire overlap between different donors. First, we examined the clonal IgG4 repertoires from a bird’s-eye view. In line with earlier findings, each serum IgG4 repertoire was dominated by a few dozen higher abundant clones, and unique for an individual donor (4 representative donors shown in Fig. 3A-D). Mass plots of all individual timepoints per donor have been uploaded to the code repository described in the code availability section. We further assessed the clonal overlap between all repertoires using an intensity-based similarity metric calculated with a matching function in which clones that fall within a given mass and retention time tolerance are considered identical (see Supplemental Figure 4). The resulting similarity matrix demonstrated only minimal inter-donor overlap between all 69 analyzed samples. In contrast, a high degree of overlap was observed in the intra-donor longitudinal samples (Fig. 3E). Strikingly, these longitudinally sampled repertoires were often stable over multiple years, as shown for donor MuSK2 where more than 50% of the detected repertoire did not change over the course of 5 years (Fig. 3F). This lack of change in the measured repertoires within a donor reveals that a large portion of high abundant IgG4 clones remain stable in sequence for several years. This finding agrees with what we earlier observed and reported for IgG1 repertoires (over one year) in healthy donors^34^, and demonstrates that there is limited ongoing somatic hypermutation in the B cells producing these antibodies.

### Detailed investigation of repertoire changes over time and treatment

The large number of longitudinal patient samples analyzed here allowed us to investigate the course of the IgG4 immune response in depth. We observed that IgG4 repertoires were unique and, in many cases, persisted for several years. At the individual level, however, each repertoire history is unique and must be interpreted alongside each patient’s clinical interventions and medications.

Figure 4 illustrates the temporal interconnection of treatments, IgG4 repertoires, and clinical scores of 6 exemplary donors, each with their own unique individual treatment regimen. Of the 6 donors, 2 (AChR24 and AChR39) received symptomatic treatment, 2 received additionally immunosuppressive treatment (MuSK5 and MuSK4), and the last 2 received immunosuppressive treatment which included rituximab (MuSK1 and MuSK7), as well as IgG modulating treatment (only MuSK1).

**Figure 4:**
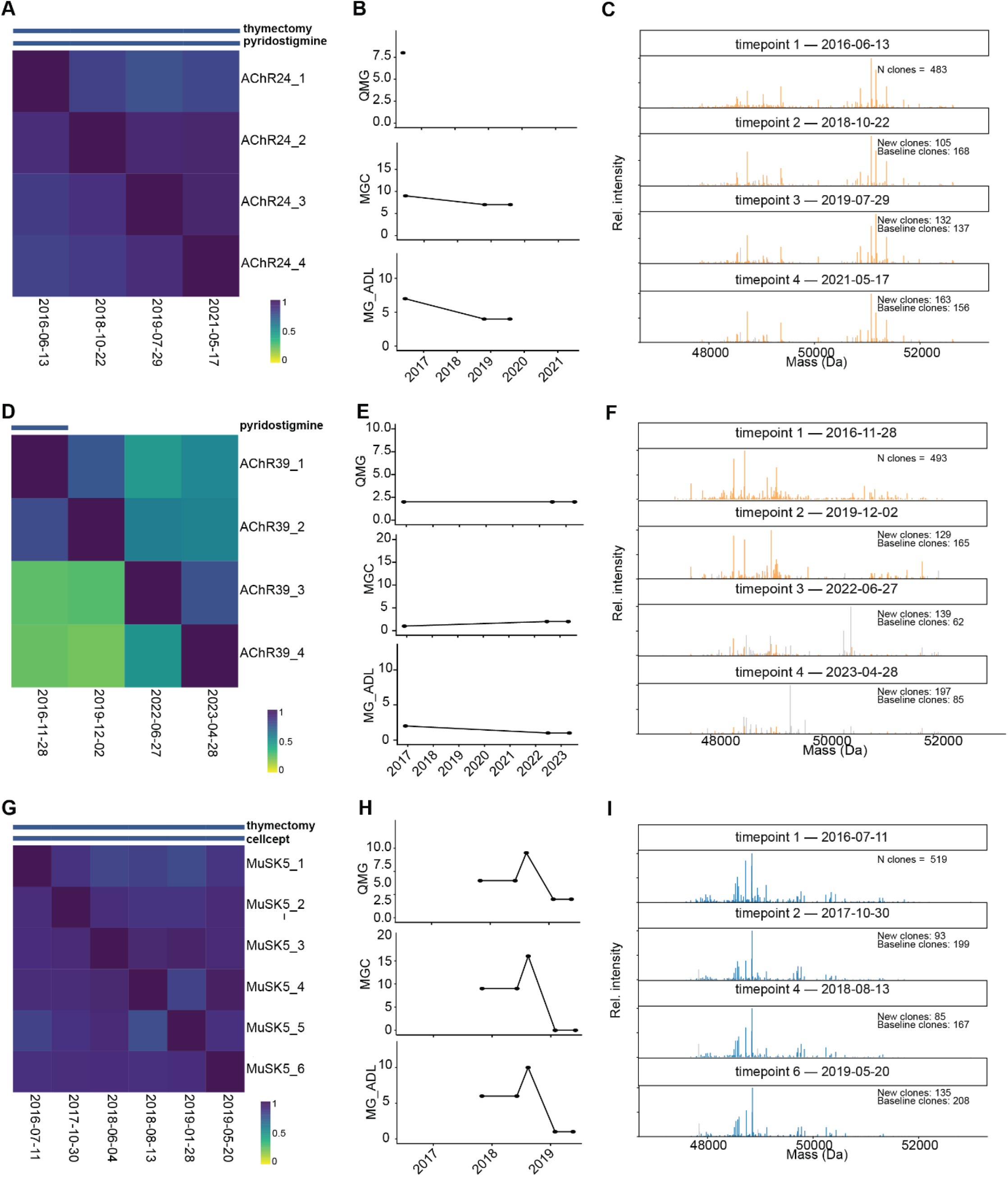
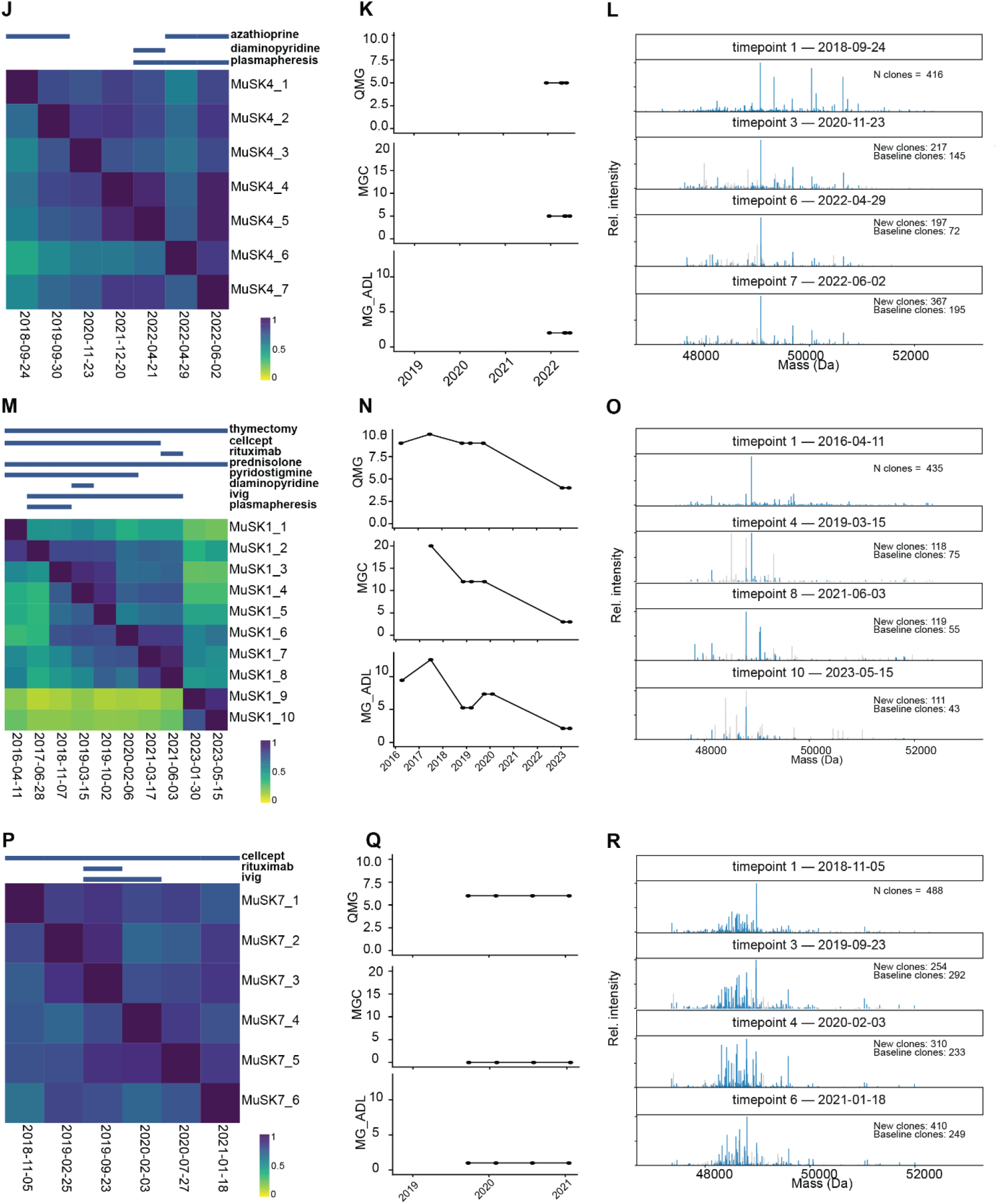
The IgG4 repertoire history is unique to each donor. Intensity-based similarity matrix annotated with ongoing treatments at sampling (blue), disease severity scores and selected mass intensity plots for two donors receiving only symptomatic treatments **(A-C, D-F)**, two donors receiving immunosuppressive treatments **(G-I, J-L)** and two donors receiving rituximab in addition to other treatments **(M-O, P-R)**. Scores in the heatmaps indicate pairwise intensity-based IgG4 repertoire similarity. These scores range between 0 (no shared clonal intensity) to 1 (all detected clonal intensity stems from clones in the respective counterpart).

Donor AChR24 represents donors whose IgG4 repertoire or disease state changed little during symptomatic treatment. The repertoire of this donor remained remarkably stable over the entire sampling duration of 5 years (Figure 4A, C). The donor had undergone thymectomy before the first timepoint and was treated throughout the study with the acetylcholine esterase inhibitor pyridostigmine. Clinical scores, available for 3 out of 4 timepoints, indicate stable disease (Figure 4B). Donor AChR39, who also received only symptomatic treatment (pyridostigmine at the first timepoint), experienced a shift in IgG4 repertoire between the first two and the last two timepoints (Figure 4 D, F)), potentially caused by immune challenges during the 2.5-year interval between timepoints 2 (December 2019) and 3 (June 2022). During the study, disease severity remained stable (Fig 4E).

For donors treated with immunosuppressants (for example MuSK5 (CellCept) and MuSK4 (azathioprine)), we observed only small repertoire changes over a 3- to 4-year interval (Figure 4G, I, J, L). During the study, disease severity for donor MuSK5 reduced markedly (Fig. 4H). The exceptional repertoire stability in both donors is striking, given that both CellCept and azathioprine act by blocking purine synthesis, thereby depleting B cells. This is particularly notable in donor MuSK5, who showed a marked reduction in disease severity despite a highly stable repertoire.

MuSK1 represents donors treated with a variety of symptomatic, immunosuppressive, and IgG modulating treatments. Over the 7 years during which the donor was studied, the repertoire underwent several major changes, including the appearance of highly abundant clones (Fig. 4M, O). The repertoire shifted markedly between time points 8 and 9, coinciding with the start of rituximab administration. During the entire follow-up period, disease activity was high but decreasing over time (Fig. 4N). Even though some repertoire changes were detected over the years, heavy IgG modulating treatments such as IVIg and plasmapheresis minimally affected the constant highly abundant, detectable IgG4 clones (Fig. 4O). Donor MuSK7 was also treated with rituximab in addition to CellCept, but the IgG4 repertoire of this donor remained largely stable over 2 years (Fig. 4P, R). Here, disease severity remained stable after treatment with rituximab (Fig. 4Q). Together, MuSK1 and MuSK7 showcase how even similar treatment regimens coincide with vastly different IgG4 repertoires and clinical trajectories.

## Discussion

Even though often considered the anti-inflammatory IgG subclass, IgG4 plays a pivotal role in some autoimmune diseases. MuSK myasthenia gravis is a prime example of such an IgG4-driven autoimmune disease. IgG4 titers are slightly elevated in MuSK MG but vary as widely as titers in healthy individuals and are therefore not suitable as biomarker^30^. In similar fashion, development of the general and IgG4-specific B-cell populations in MuSK MG patients is comparable to that of healthy individuals, both in terms of maturity and plasma cell proportions^35^. We therefore investigated whether abnormalities could be found on the protein level of the individual serum IgG4 antibody repertoire. As controls we included sera from donors with other myasthenic disorders such as AChR MG and LEMS, as well as healthy donor sera. We processed a uniquely large cohort of 218 samples obtained from 69 donors, making this study the largest clonal profiling investigation yet.

In line with previous observations on antibody repertoires in IgG1^28^, IgA1^29^, and IgG4^10^, we observed for each of the 69 donors that their respective IgG4 repertoire functions as a unique clonal fingerprint. Furthermore, within this cohort we were able to study much more extended time courses than in our previous studies. Strikingly, our conclusions on repertoire stability here (> 5 years) are not different from the one-year timeline: most clones do not decay. Thus, extending previous observations, we show repertoire stability over a period of more than 5 years (e.g., donor MuSK2 showing 44% repertoire stability over 8 years). This aligns with previous assumptions that large parts of the repertoire are secreted by long-lived plasma cells (LLPCs)^34^. Moreover, even treatment of AIDs aimed at modulating B cell populations do not seem to affect the repertoire as we observe it here. The production of a large fraction of the repertoire by LLPCs may explain the lack of effect of rituximab on IgG4 repertoires in some patients, since CD20 expression in LLPCs is low to absent^36-38^. Consistent with this, we observed close to no effect of the B cell depleting (but LLPC sparing) CellCept or azathioprine treatments on IgG4 repertoires. This dichotomy of a stable repertoire despite rituximab’s high clinical efficacy in MuSK MG^39, 40^ suggests two compartments from which IgG4 responses originate: a population of LLPCs generating a highly abundant and stable IgG4 repertoire, and a population of short-lived plasma cells that, in MuSK MG, generates lowly abundant autoreactive IgG4 clones. This hypothesis is in line with earlier observations of decreasing anti-MuSK titers combined with stable total IgG4 titers after rituximab administration^41^. It is further supported by donors in this study who showed strong clinical improvements while being treated, despite a highly stable IgG4 repertoire (e.g., donor MuSK5; Fig. 4G-I). Considering a previous report on the lower abundance of IgG4 B cells in spleen and lymph nodes, it is tempting to speculate that IgG4 LLPCs reside in the bone marrow where they may be the source for the overall relatively stable IgG4 repertoire^42^.

An important and unique feature of IgG4 is FAE. Our analysis across 218 samples revealed that in serum about 22% of IgG4 are not held together by disulfide bridges but exclusively by non-covalent interactions, i.e. they are available for exchange. All deviations fell into the range observed earlier in another cohort of healthy donors^10^, indicating that IgG4 half-molecule dissociation can occur to a similar extent across all studied groups. We further observed an increased number of detected IgG4 clones for MuSK patients compared to healthy donors in the baseline sampling points. This is opposite of our initial hypothesis that a dominant IgG4 AID could drive the outgrowth of autoreactive clones and therefore present with lower clonal numbers. The increased number of clones in MuSK MG patients, however, also coincide with a general tendency for slightly elevated IgG4 titers in MuSK MG patients^30^, and may indicate an overall inflated repertoire. Another explanation may be found in the increased levels of (anti-MuSK-)IgG4 Fab glycosylation^43, 44^. The same clone may, for example, carry different glycans and consequently present as a larger number of clones. However, we did not find any proof of this in our data. Lastly, at later time points, the increased number of IgG4 clones in MuSK MG is still visible, although not statistically significant. This may be the result of treatments the patients received over time, as all treatments correlated with lower clone numbers. Since most treatment regimens combine different modalities, we cannot conclude which specific treatment decreased the detected clone number in these cases.

We next investigated whether clone numbers correlated with clinical disease scores and found a negative correlation between IgG4 clonality and two patient-reported disease scoring instruments (MG-ADL and MGC) in MuSK MG patients. Additionally, we observed no strong treatment effects on repertoire composition that would explain this (see below). In AChR MG patients, total IgG4 clone numbers correlated with anti-AChR autoantibody titers, which is striking given that AChR MG is IgG1/IgG3-mediated. This relationship was absent in LEMS patients (also IgG1/IgG3-mediated) and in MuSK MG patients (IgG4-mediated). Anti-MuSK titers in MuSK MG did not correlate with total IgG4 clone numbers, indicating that there likely is no expansion of highly abundant MuSK-specific IgG4 clones. This is in line with anti-MuSK titers generally making up just a small fraction of total IgG4 titers^41^.

Taken together, our longitudinal dataset reveals that IgG4 repertoires are unexpectedly stable over time in both autoimmune patients and healthy controls, even in the face of strong immunosuppressive or immunoglobulin-modulating treatments. More specific for the AIDs in this study, our findings show that the number and distribution of individual clones in the repertoires of MuSK MG patients is normal, suggesting that MuSK MG pathology is driven by a low abundance, polyclonal anti-MuSK response. Expanding the repertoire profiling method to identify antigen-specific clones is required to provide insight into the autoreactive IgG4 repertoire. This could be particularly interesting for monitoring clinical success of new chimeric autoantibody receptor T cell^41^ therapies.

## Materials and Methods

### Origin of the serum samples and recombinant mAbs

Individual serum samples were collected anonymously in accordance with the ethics board of the Leiden University Medical Center (LUMC, Leiden, the Netherlands) under CME protocol number P17.011. All subjects provided written informed consent prior to participation, and experiments were in accordance with the Declaration of Helsinki, including current revisions, and Good Clinical Practice guidelines. The samples were stored at ™80°C until used for processing. For method standardization we used the IgG4 monoclonal antibody Natalizumab which was purchased from Evidentic (Potsdam, Germany).

### Serum IgG4 purification and digest

To capture IgG4 from serum, the procedure was executed as described previously^10^. In brief, 20 µL CaptureSelect IgG4 (Hu) affinity matrix (Thermo Fisher Scientific, Leiden, Netherlands) slurry was added to Pierce spin columns with screw cap (Thermo Fisher Scientific, Rockford, Illinois, USA). The slurry was washed three times by adding 150 µL anticoagulant (375 mM saccharose, 82.5 mM Sodium citrate, 17.5 mM citric acid monohydrate), followed by centrifugation at 500 x g and room temperature (RT). A plug was inserted to the bottom of the spin column, and 100 µL human serum spiked with 200 ng Natalizumab IgG4 was mixed with 50 µL anticoagulant. The samples were incubated on an Eppendorf thermal shaker (Wesseling-Berzdorf, Germany) with a ThermoTop for 1h, 750 rpm at RT. Following incubation, the plugs were removed from the spin column and the IgG4 beads were washed three times with 200 µL PBS (+ 1 M NaCl) and two times with 200 µL MilliQ ddH_2_O. Purified IgG4 were eluted from the matrix into 12 µL phosphate neutralization buffer (429.3 mM sodium dihydrogen phosphate monohydrate, 470.7 disodium hydrogen phosphate dihydrate, pH 7.54) in LoBind Eppendorf tubes in three elution steps. Each time, 20 µL 0.1 M glycine (pH 2.7) were added to the IgG4 beads. After incubation for 5 min, 750 rpm at RT on an Eppendorf thermal shaker, the spin columns were centrifuged at 1000 x g. For hinge-direction digestion, 2.5 µg FabRICATOR (IdeS; Genovis, Kävlinge, Sweden) were added in 50 uL PB to the sample and incubated for 18 h at 750 rpm and 37 °C on an Eppendorf thermal shaker with a ThermoTop. Afterwards, Fc and undigested antibodies were depleted from the sample by repeating the IgG4 enrichment described above. Subsequently, the spin columns were centrifuged for 1 min at 1000 x g and IgG4 Fab and Fab2 were collected in LoBind Eppendorf tubes.

### Size exclusion chromatography (SEC)

Size exclusion chromatography was performed on an Agilent 1260 Infinity HPLC system (Agilent, Santa Clara, California, USA) operated in isocratic flow with an attached SEC column (ACQUITY UPLC BEH SEC, Waters, Framingham, Massachusetts, USA, 200 Å, 1.7 µm particle size, 4.6 mm X 300 mm). A 20-minute separation method was applied (100 mM ammonium acetate, pH 6.9) at a flowrate of 200 µL*min-1 for separation. Fabs were collected in one fraction between 13.8–16.0 minutes. Collected samples were snap frozen in liquid nitrogen and subsequently lyophilized (Labconco Freezone 2.5 L -80°C, Kansas City, Missouri, USA). Prior to further analysis, each lyophilized sample was reconstituted in PBS (137 mM sodium chloride, 2.7 mM potassium chloride, 4.77 mM sodium dihydrogen phosphate monohydrate, 5.23 mM disodium hydrogen phosphate dihydrate) in a volume adjusted to the integrated UB absorbance of the corresponding SEC trace.

### LC-MS data acquisition

Intact Fab samples were analyzed by LC-MS using a Vanquish Neo UHPLC (Thermo Fisher Scientific, San José, California, USA) coupled to an Orbitrap Exploris 480 (Thermo Fisher Scientific, Bremen, Germany). Per injection, 500 ng of total protein dissolved in PBS was loaded onto a MabPac RP analytical column (Thermo Fisher Scientific, San José, California, USA, 150 µm x 150 mm, 4 µm particle size), with protein concentrations quantified beforehand on a NanoDrop spectrophotometer. Chromatographic separation was performed by applying a 54-minute gradient from 28% to 38% solvent B (0.1% FA, 0.02% DFA in ACN) at a flow rate of 150 μL*min^-1^ to the mobile phase (0.1% FA, 0.02% DFA in ddH_2_O). MS data acquisition was performed in Intact Protein and Low Pressure mode with positive ionization. Source parameters were as follows: Spray voltage was set at 2 kV, capillary temperature to 275 °C, and probe heater temperature to 100 °C. The RF Lens was set to 60% and source fragmentation was activated at 15 V. An MS1 scan between 800 – 3.200 m/z was applied with a maximum injection time (maxIT) of 50 ms, a resolution of 7500, and a normalized automatic gain control (AGC) target value of 300%.

### Clonal profiling data processing and analysis

Deconvolution of RAW files was carried out in BioPharmaFinder 3.2 (Thermo Fisher Scientific, San José, California, USA) using the ReSpect algorithm. Source spectra were generated by the sliding windows algorithm across a retention time between 6 and 60 minutes with the following parameters: 0.1 min target average spectrum width, 25% offset, 30 ppm merge tolerance, and 95% noise rejection. The deconvolution output was constrained to a mass range between 20 000 to 65 000 Da, target mass was set to 48 000 Da and charge states between 10 and 60 were included. Output files were further analyzed using Python 3.10.9 (libraries: Pandas 1.5.3^45^, Numpy 1.23.5^46^, Scipy 1.12.0^47^, Matplotlib 3.7.0^48^, and Seaborn 0.12.2^49^). Component masses were recalculated using an intensity weighted mean considering only the most intense peaks comprising 90% of the total intensity. Components between 46 and 55 kDa with the most intense charge state above 1,000 Thomson and a score above 40 were considered as derivatives of IgG4 clones. The clones were matched between runs using average linkage (unweighted pair group method with arithmetic mean UPGMA) L^∞^ distance hierarchical clustering. Flat clusters were formed based on a cophenetic distance constraint derived from the mass and retention time tolerance. These tolerances were defined as 1.5 Da and 3 minutes for Fab. Clones within a flat cluster were considered identical between runs. Retention times were aligned based on the spiked in Natalizumab to minimize deviations between runs. To exclude shared background contaminants such as Fc fragments or co-purified serum proteins, components detected in more than 15 of the 69 donors were discarded from further analysis.

### Serum titer analysis

Anti-LRP4, anti-VGCC and anti-AChR antibody serum titers were determined by the Clinical Chemical Lab of the LUMC using their standardized and validated ELISA assays. Anti-MuSK IgG4 titers were determined by an ELISA assay developed in-house at the Department of Human Genetics and Department of Neurology of the LUMC, as previously described^50^.

### Statistical analysis

Statistical analysis and data visualization were performed using R 4.3.1. Statistical significance was assessed by linear mixed effects models using the *nlme* R package version 3.1-168 or by analysis of variance models using the *stats* core R package version 4.3.3 followed by Tukey Honest Significant Differences post-hoc analysis using the *rstatix* R package version 0.7.3^51^. Linear regression analyses were performed using the *ggpmisc* R package version 1.0.0^52^.

## Conflict of interest

MGH and JV are coinventors on patent applications based on MuSK-related research. The LUMC receives royalties from IBL and Argenx, MGH and JV receive royalties from Argenx. AJRH is co-founder of Abvion. All other authors report no disclosures.

### Acknowledgements

We gratefully acknowledge the patients included in this study, as well as the LUMC Vrijwillige Donoren Service (LuVDS) and its volunteers for providing donor materials. We thank prof. dr. Silvère van der Maarel for his advice and support. We would further like to express our gratitude to Anna van Doorn and Juliëtte Lenssen for their effort in collecting clinical data. The LUMC is part of the European Reference Network for Rare Neuromuscular Diseases (ERN EURO-NMD) and the Netherlands Neuromuscular Disorders Center (NL-NMD).

## Author contributions

**Linus Wollenweber:** methodology, investigation, software, formal analysis, writing – original draft, data curation, visualization

**Laurent M. Paardekooper:** investigation, software, formal analysis, writing – original draft, data curation, visualization

**Yvonne J.M. Campman:** methodology, data curation

**Jessica M. van Bokkum:** methodology

**Arjan Barendregt**: resources, supervision

**Martijn R. Tannemaat:** resources, supervision

**Jan J.G.M. Verschuuren:** resources, supervision

**Albert J.R. Heck:** conceptualization, resources, supervision, funding, writing – review & editing

**Albert Bondt:** conceptualization, resources, supervision, writing – review & editing

**Maartje G. Huijbers:** conceptualization, resources, supervision, writing – review & editing

