## Supplemental Figures and Tables for "IgG4 antibody repertoire diversity is stable over several years in health and autoimmune disease"

**Contains Supplemental Figures 1, 2, 3, 4, and Supplemental Table 1, 2, 3**

**A**

**B**

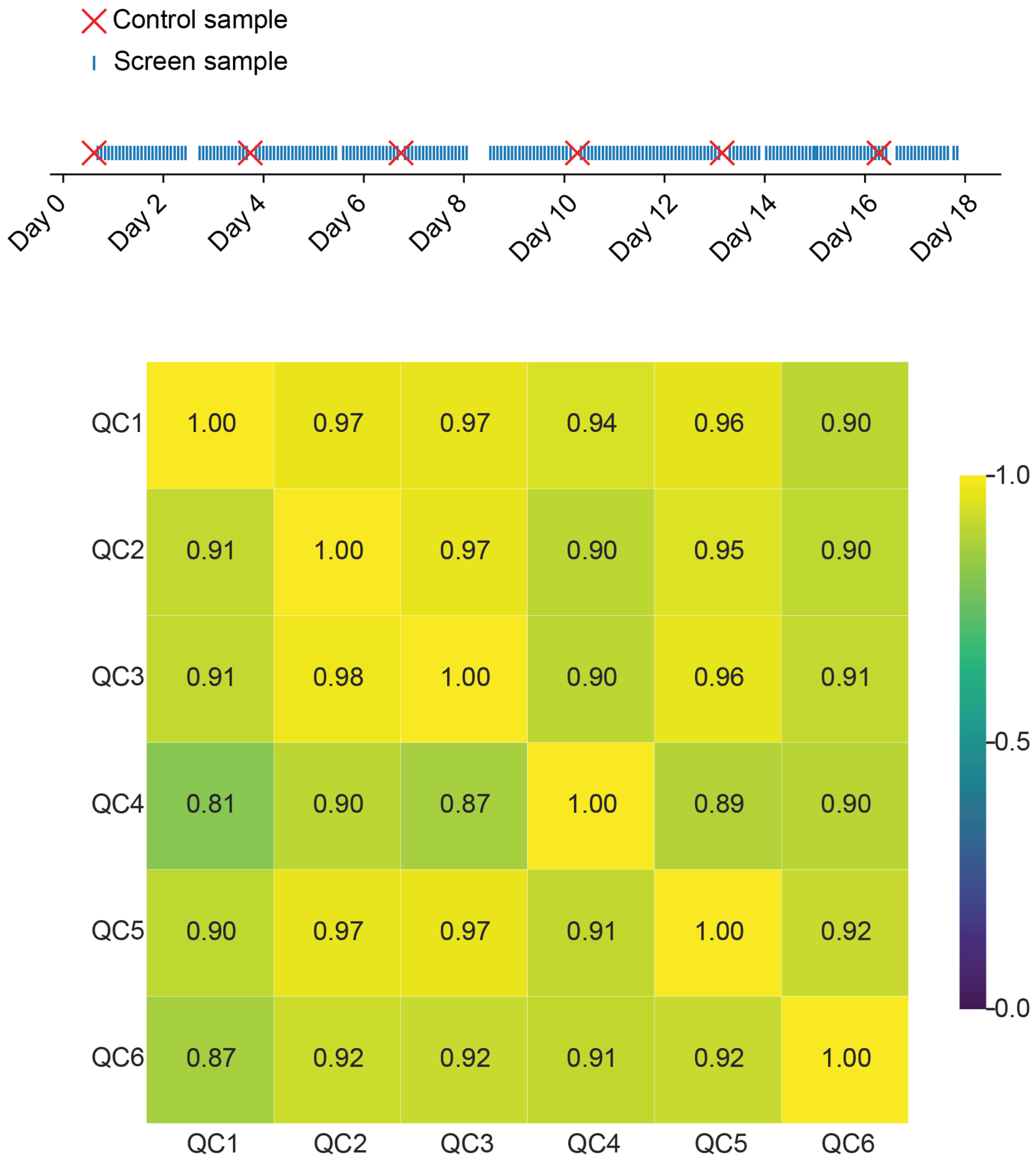


**Figure S1 | Repeated measurements of a control sample over time.** *A control IgG4 Fab sample was repeatedly measured over the 18 days days of mass spectrometric data acquisition (A). The chart indicates screen sample measurements (blue lines) and regularly interspersed control measurements (red crosses). The constant performance of the LC-MS system can be proxied by the similarity of the QC samples over time (B). The similarity, based on the intensity-based overlap of the deconvoluted repertoire is indicated by the number in each field using the ruler on the right. The proportion of the intensity of clones identified within each pairwise comparison is assigned a value between 0 (no shared intensity) and 1 (all intensity can be explained by clones found in the respective counterpart).*

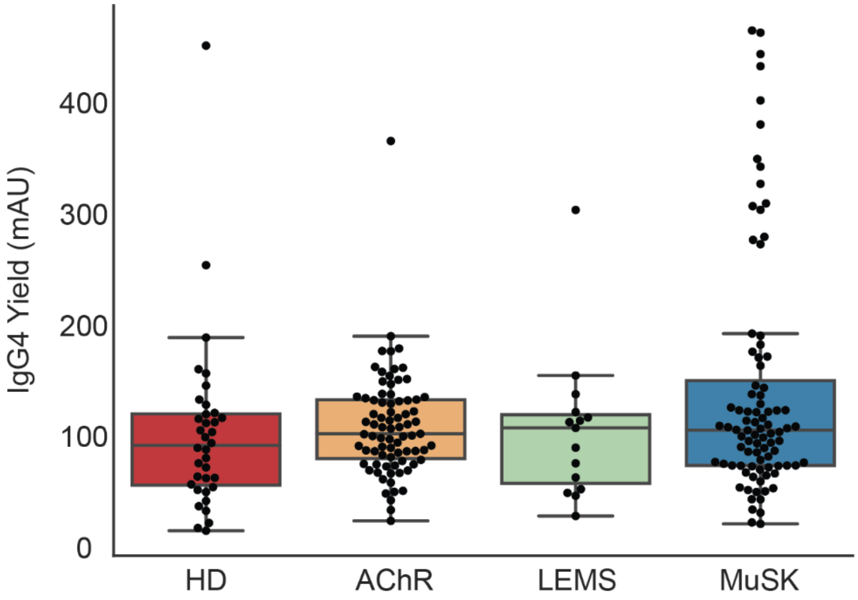


**Figure S2 | IgG4 fragment yield after binding and digestion.** *Fab2 and Fab yield were quantified for each sample by integrating SEC-UV peak areas between 11.6 and 16 minutes and are stated as arbitrary units (mAU). Each boxplot displays median, and interquartile range (IQR; box edges = 25$^{th}$ and 75$^{th}$ percentile). Whiskers extend to the most extreme datapoint within 1.5 x IQR from the box edges; outliers are plotted as diamonds. Each donor group is shown separately. No statistical difference was found between the groups with p < 0.5 in unpaired Student's t-tests.*

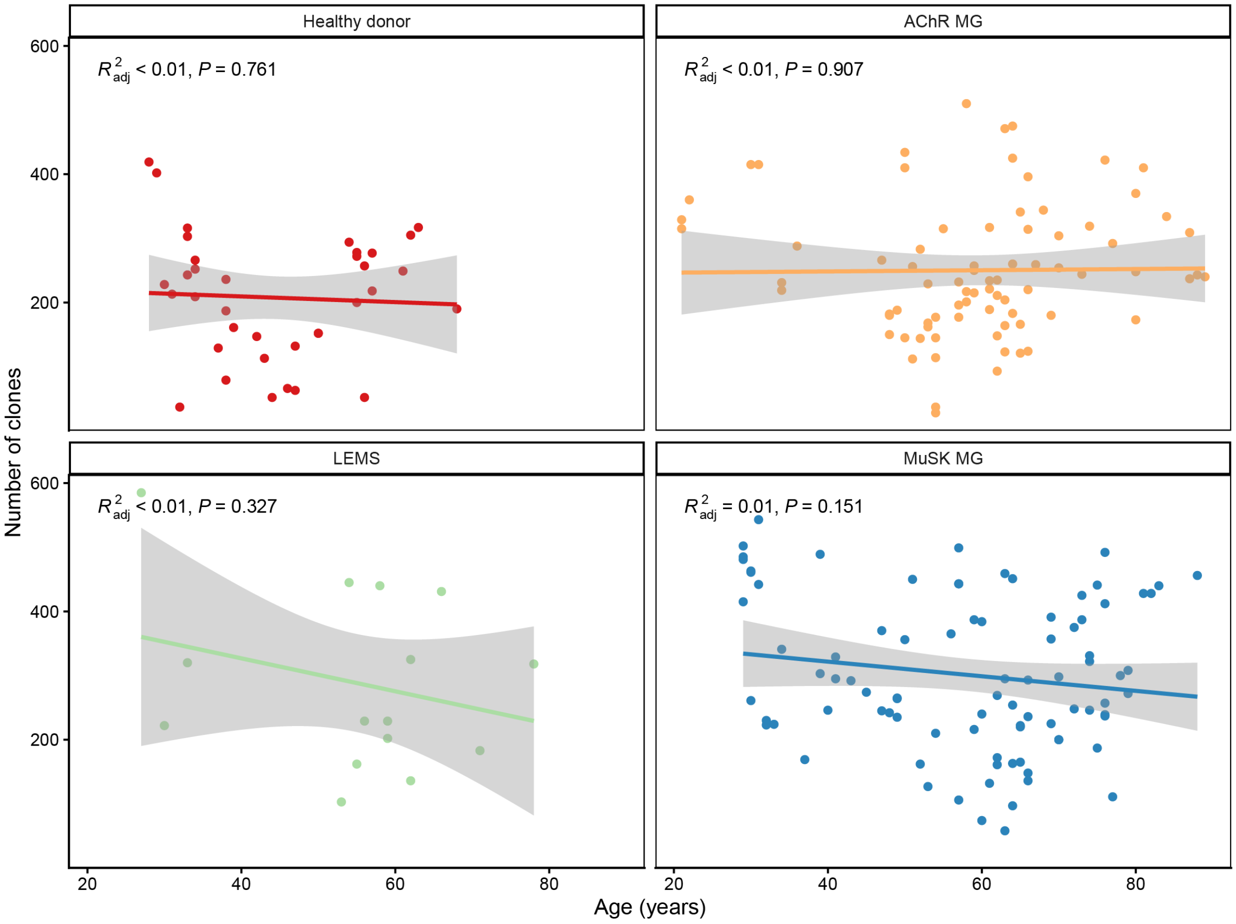


**Figure S3 | Relation between number of detected clones and donor age.** *Scatter plot of IgG4 clone count versus donor age at blood withdrawal. Line shows linear regression with 95% confidence interval (shaded area). $R^2$ and p-values are indicated in the figure.*

**Example 1:** Two highly similar samples are compared. Depending on the intensity of not-shared clones, the similarity score differs and is unique for each sample within a comparison.

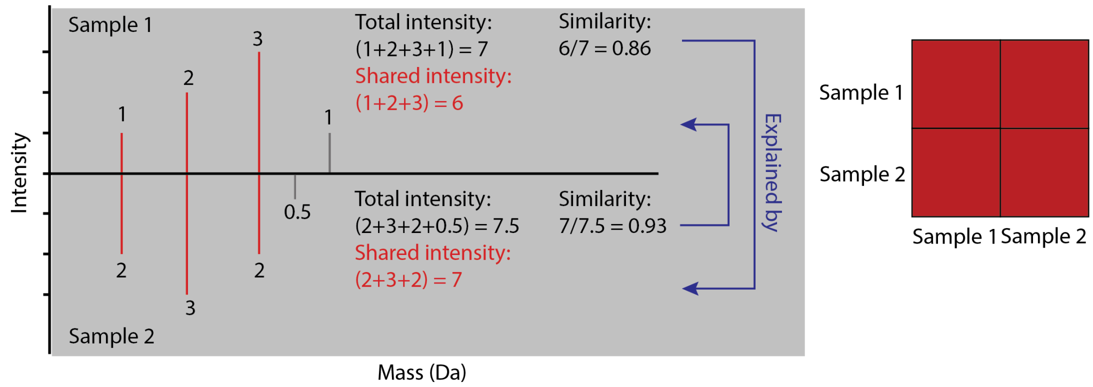


**Example 2:** Two mostly different samples are compared. Since sample 2 has only few high abundant clones, one high abundant shared clone can skew the similarity score to a one-sided high similarity.

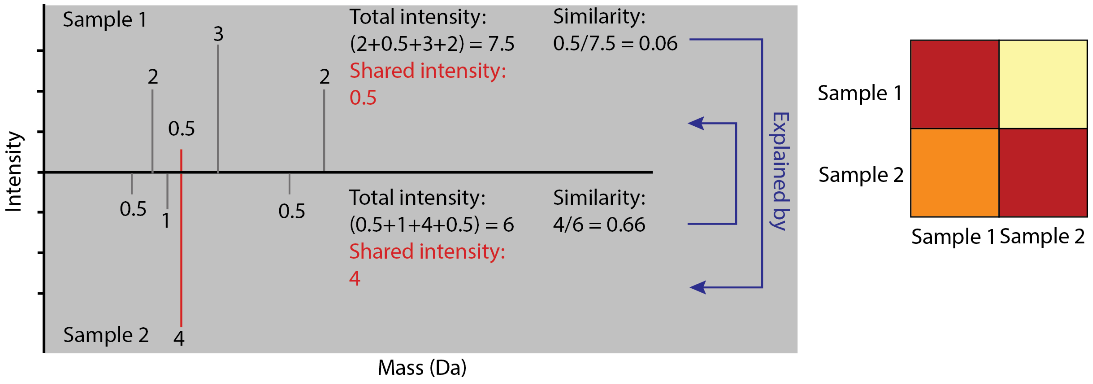


**Figure S4 | Calculating clonal similarity.** *Components identified in BioPharmaFinder receive unique IDs based on hierarchical clustering. In pairwise comparisons, the proportion of the total sample intensity that can be explained by a respective counterpart is given as a value between 0 and 1, - the similarity.*

**Table S1 | Donor demographics.** *List of donor demographics, sorted by group (healthy donor, AChR MG, LEMS, MuSK MG).*

| | **Healthy donor** N = 36[1] | **AChR MG** N = 79[1] | **LEMS** N = 15[1] | **MuSK MG** N = 88[1] | **p-value**[2] | **q-value**[3] |
|---|---|---|---|---|---|---|
| **Demographics** | | | | | | |
| Sex | | | | | 0.2 | 0.3 |
| *F* | 18 (50%) | 44 (56%) | 11 (73%) | 41 (47%) | | |
| *M* | 18 (50%) | 35 (44%) | 4 (27%) | 47 (53%) | | |
| Age at sampling (years) | 44 (34, 55) | 61 (52, 66) | 58 (53, 62) | 62 (47, 71) | <0.001 | <0.001 |
| Number of clones | 216 (140, 275) | 235 (177, 315) | 229 (183, 431) | 283 (223, 414) | <0.001 | 0.002 |
| Clonal dispersity (range: 0-1) | 0.38 (0.32, 0.48) | 0.40 (0.32, 0.48) | 0.41 (0.33, 0.57) | 0.39 (0.30, 0.48) | 0.5 | 0.5 |
| **Diseases characteristics** | | | | | | |
| Age at diagnosis (years) | — | 51 (36, 62) | 51 (26, 54) | 43 (26, 52) | 0.016 | 0.039 |
| Disease duration at sampling (years) | — | 8 (4, 14) | 8 (4, 25) | 12 (7, 20) | 0.010 | 0.039 |
| MG-ADL | — | 5.0 (3.0, 8.0) | 4.0 (4.0, 8.0) | 3.0 (1.0, 6.0) | 0.2 | 0.3 |
| QMG | — | 5.5 (4.0, 8.0) | 8.0 (1.0, 10.0) | 5.0 (3.0, 7.0) | 0.6 | 0.6 |
| MGC | — | 7.0 (5.0, 12.0) | 7.0 (0.0, 7.0) | 5.0 (1.0, 9.0) | 0.093 | 0.2 |
| MGFA | — | | | | | |
| *0* | — | 33 (43%) | 0 (0%) | 43 (51%) | | |
| *1* | — | 25 (32%) | 0 (0%) | 3 (3.5%) | | |
| *2* | — | 12 (16%) | 3 (100%) | 12 (14%) | | |
| *2a* | — | 4 (5.2%) | 0 (0%) | 0 (0%) | | |
| *2b* | — | 2 (2.6%) | 0 (0%) | 16 (19%) | | |
| *3* | — | 0 (0%) | 0 (0%) | 2 (2.4%) | | |
| *3b* | — | 1 (1.3%) | 0 (0%) | 9 (11%) | | |

| **Treatments** | | | | | | |
|---|---|---|---|---|---|---|
| Immunosuppression naive at sampling | — | 21 (27%) | 3 (20%) | 1 (1.1%) | <0.001 | <0.001 |
| Plasmapheresis | — | 0 (0%) | 0 (0%) | 6 (6.8%) | 0.042 | 0.066 |
| IVIg | — | 7 (9.1%) | 3 (20%) | 10 (11%) | 0.4 | 0.5 |
| Amifampridine | — | 0 (0%) | 9 (60%) | 0 (0%) | <0.001 | <0.001 |
| Diaminopyridine | — | 0 (0%) | 0 (0%) | 9 (10%) | 0.007 | 0.012 |
| Pyridostigmine | — | 45 (57%) | 5 (33%) | 14 (16%) | <0.001 | <0.001 |
| Prednisolone | — | 24 (30%) | 6 (40%) | 40 (45%) | 0.14 | 0.2 |
| Azathioprine | — | 27 (34%) | 1 (6.7%) | 24 (27%) | 0.082 | 0.11 |
| Rituximab | — | 1 (1.3%) | 0 (0%) | 4 (4.5%) | 0.6 | 0.6 |
| CellCept | — | 12 (15%) | 0 (0%) | 36 (41%) | <0.001 | <0.001 |
| Thymectomy | — | 40 (51%) | 0 (0%) | 26 (30%) | <0.001 | <0.001 |

[1]n (%); Median (Q1, Q3)

[2]Pearson's Chi-squared test; Kruskal-Wallis rank sum test

[3]False discovery rate correction for multiple testing

**Table S2 | Treatment categories per patient group**

| ***Treatment at sampling*** | ***Healthy donor*** *N = 36*[1] | ***AChR MG*** *N = 79*[1] | ***LEMS*** *N = 15*[1] | ***MuSK MG*** *N = 88*[1] | ***p-value***[2] | ***q-value***[3] |
|---|---|---|---|---|---|---|
| *Immunosuppression* | *0 (NA%)* | *46 (58%)* | *7 (47%)* | *71 (81%)* | *0.001* | *0.002* |
| *IgG modulation* | *0 (NA%)* | *7 (9.1%)* | *3 (20%)* | *14 (16%)* | *0.3* | *0.3* |
| *Symptomatic* | *0 (NA%)* | *45 (57%)* | *11 (73%)* | *22 (25%)* | *<0.001* | *<0.001* |

[1]*n (%)*

[2]*Fisher's exact test*

[3]*False discovery rate correction for multiple testing*

**Table S3 | Sequence of the spiked-in monoclonal antibody.**

| mAb | Heavy chain (Hc) | Light chain (Lc) | Fab mass<br>Fd mass<br>Lc mass | Modification |
|---|---|---|---|---|
| Natalizu-mab | QVQLVQSGAEVKKPGASVKVSCKASGF<br>NIKDTYIHWVRQAPGQRLEWMGRIDPA<br>NGYTKYDPKFQGRVTITADTSASTAYME<br>LSSLRSEDTAVYYCAREGYYGNYGVYA<br>MDYWGQGTLVTVSSASTKGPSVFPLAP<br>CSRSTSESTAALGCLVKDYFPEPVTVSW<br>NSGALTSGVHTFPAVLQSSGLYSLSSVV<br>TVPSSSLGTKTYTCNVDHKPSNTKVDKR<br>VESKYGPPCPSCPAPEFLGGPSVFLFPP<br>KPKDTLMISRTPEVTCVVVDVSQEDPEV<br>QFNWYVDGVEVHNAKTKPREEQFNST<br>YRVVSVLTVLHQDWLNGKEYKCKVSNK<br>GLPSSIEKTISKAKGQPREPQVYTLPPSQ<br>EEMTKNQVSLTCLVKGFYPSDIAVEWES<br>NGQPENNYKTTPPVLDSDGSFFLYSRLT<br>VDKSRWQEGNVFSCSVMHEALHNHYT<br>QKSLSLSLGK | DIQMTQSPSSLSASVGDRVTITCKTSQDI<br>NKYMAWYQQTPGKAPRLLIHYTSALQPG<br>IPSRFSGSGSGRDYTFTISSLQPEDIATYY<br>CLQYDNLWTFGQGTKVEIKRTVAAPSVFI<br>FPPSDEQLKSGTASVVCLLNNFYPREAK<br>VQWKVDNALQSGNSQESVTEQDSKDST<br>YSLSSTLTLSKADYEKHKVYACEVTHQGL<br>SSPVTKSFNRGEC | 49,209.0 Da<br>25,664.8 Da<br>23,556.3 Da | Pyrogluta-mate (Hc) |
